# A hierarchy of locomotion costs shapes optimal foraging strategy

**DOI:** 10.64898/2026.04.14.718484

**Authors:** Thomas P. Ilett, Omer Yuval, Felix Salfelder, Robert I. Holbrook, David C. Hogg, Thomas Ranner, Netta Cohen

## Abstract

Animals must make decisions about how to move through their environment to find food, shelter and mates, yet the general principles underpinning effective search strategies remain poorly understood. Exploration patterns, such as alternating intensive local search with long-range relocation, are observed across many species, but whether these reflect universal organising principles, or are simply imposed by structured environments, is an open question. Here, we reconstruct hours of *Caenorhabditis elegans* foraging in homogeneous three-dimensional volumes and show that this animal achieves optimal volumetric coverage using a hierarchical search strategy of quasi-planar locomotion patches punctuated by costlier volumetric reorientations. We propose that underlying this cost hierarchy is the worm’s highly-constrained shape-space and show that despite being spanned by only five body-posture modes, it gives rise to effective locomotion, combining non-planar gaits, planar turns and three-dimensional reorientation manoeuvres. Our finding that action frequencies decay with inferred cost in a Boltzmann-like distribution is consistent with the Principle of Maximum Entropy: optimal search emerging from the tendency to maximise information gained subject to embodied constraints. This general principle offers a unifying framework to assess the optimality of animal decision-making, with implications for understanding nervous system function, foraging ecology, and the physics of living active matter.

## Introduction

Animals perform local search to explore their environments. Common strategies for search are often found across species despite vastly different body geometries, gaits and environments. Typically, animals perform local search using bouts of forward motion, punctuated by pauses (also called intermittent locomotion^1^) and turning events.^2,3^ Randomly sampling forward bouts and turns results in diffusive or super-diffusive spatial exploration,^4^ whereas biasing turn probabilities and turn angles can generate effective navigation towards preferred targets.^2,5^ Decomposing exploration trajectories into action primitives such as locomotion gaits and reorientations provides a useful framework for identifying behavioural choices, their costs and constraints, but risks yielding species-specific insights that fail to generalise. However, a decomposition in terms of a common currency – the relative costs of different actions – should reveal unifying principles that inform the spatial exploration strategy used by different animals to achieve their goals.^6^

Whether on a substrate or in 3D environments, and especially in heterogeneous or structured environments, exploration strategies often manifest in search spaces composed of local patches.^7–11^ In addition, most animals exhibit a strong directional bias due to gravity, light or physical structures of the local environment. To reveal innate 3D strategies, we asked how animals behave in uniform volumes. In the absence of directional cues, any directional bias would be dominated by the embodied nature of the search. The widely studied nematode *C. elegans* is an ideal model system for linking locomotion to exploration strategies. While it inhabits complex fluid, soft matter and granular environments in nature,^12,13^ in 3D, neither its modes of locomotion nor its exploration strategies are known. With some exceptions,^14–18^ nematode locomotion is described in terms of planar sinusoidal undulations.^14,19^ However, to survive in their natural environment – decaying vegetation^12,13,20^ – they must navigate a volume; too close to the surface they become vulnerable to predators, too deep and food becomes scarce.^21,22^ In uniform 3D environments and on the scale of a rotting apple (or up to hundreds of body lengths) though, one might expect any directional bias to disappear and exploration to be fully isotropic.

*C. elegans* sinusoidal undulations are generated by propagating a smooth curvature wave along the body,^14,23–26^ which is explained by their planar neuromuscular connectivity pattern below the neck.^27,28^ A more complex neural circuit innervating muscles in the head and neck enables greater freedom of movement that is also observed through head lifting on a substrate.^19^ Previous studies of 3D locomotion in *C. elegans* adults have reported slight deviations from planar undulations,^29^ a novel three-dimensional roll manoeuvre,^16^ the first reported observation of a three-dimensional trajectory of an animal immersed in a volume^17^ and observations of animals traversing barriers on non-planar substrates.^30^ In addition to adults, *C. elegans* dauer larvae display complex 3D individual^31^ and collective behaviours.^32,33^

### *C. elegans* explore 3D space using non-planar gaits and manoeuvres

To investigate the connection between *C. elegans* locomotion and local search in 3D, we designed a triaxial imaging system that enables recording of a sufficiently large volume of view (10-20 worm lengths per side) at sufficient magnification and resolution for accurate 3D posture reconstruction (Fig. 1a, Methods). We recorded nearly 7 hours of footage of individual young adult wild-type *C. elegans* across a range of non-Newtonian fluids (1-4% gelatin in buffer solution, 4-25 minutes per movie, Methods), from which we obtained a corpus of single-animal postures and trajectories (Fig. 1). This corpus reveals a wide variety of locomotor behaviours, including novel 3D gaits and complex 3D manoeuvres, which we used to characterise 3D exploration (Fig. 1b-e, Supp. Videos 1-3).

**Figure 1:**
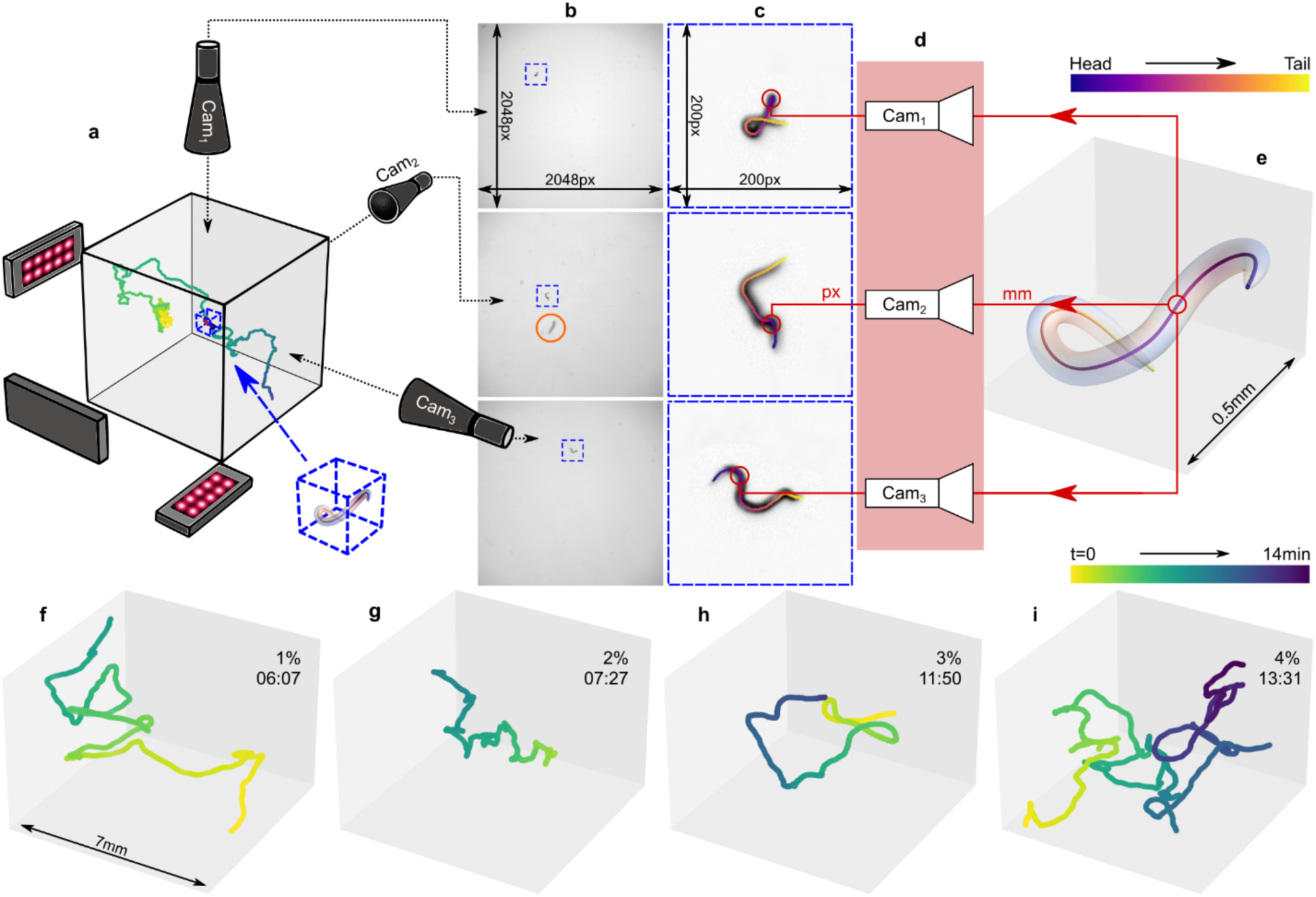
Experimental setup, tracking and 3D reconstruction pipeline. a. Setup schematic (not to scale): 10-minute reconstructed trajectory with a worm (inside blue cube). Three static telecentric lenses face the sides of a sample cube, uniformly backlit with red LED lights. b. Raw image triplet. The same worm is identified in each view (blue squares). Worms appearing in only one view (orange circle) are ignored. c. Cropped, pre-processed images. d. Pinhole camera models (Cam_1_-Cam_3_) translate between lab coordinates (mm) and image coordinates (pixels). Reconstructed midline points are projected through each camera model to the respective 2D images (red arrows). e. Reconstructed 3D posture. Projections of the 3D reconstructed postures superimposed in c (colour). f-i. Centre-of-mass trajectories (upper right: trajectory durations; scale bar in f) for different animals in 1%-4% gelatin reveal volumetric foraging trajectories, from dense searches of one or two regions to long meandering trajectories.

After removal from food on a substrate, *C. elegans* intersperse periods of forward undulations with turns^2,19,34^ to perform a local area search.^5^ Similarly to planar local area search,^2,5,19^ we find that 3D exploration consists of periods of forward locomotion subject to gentle steering and occasional reversals (backward locomotion) coupled to turn manoeuvres (Supp. Video 1). However, in 3D, *C. elegans* produce extensive volumetric exploration of the environment when moving in a variety of fluids (Fig. 1f-i). To quantify the 3D shape-space of *C. elegans*, we extracted a database of over 415,000 postures from our movies, recorded across all concentrations, and constructed an eigenbasis of these postures (Methods, Fig. 2a,e). Principal component analysis (PCA) decomposition of *C. elegans* postures on an agar surface reveals a low-dimensional shape-space, with four components (or “eigenworms”) describing 95% of the variance.^23^ Five 3D eigenworms are sufficient to capture 96% of the variance in our data, dropping to 87% for four modes (Methods, Fig. 2a,f, Ext. Fig. 2). This result establishes a low-dimensional posture space that is available to the animal at feasible cost. Similarly to 2D, 3D forward locomotion is described by oscillating contributions from the two dominant (lowest-order) eigenworms, as is backward locomotion, albeit with an opposite phase velocity (Fig. 2b,d, Ext. Fig. 3); higher-order eigenworm contributions peak during turns (Fig. 2c,d). The dominance of the first two eigenmodes strongly suggests that these bouts of forward and backward locomotion are more efficient or less costly than turning manoeuvres. In 2D, worms smoothly modulate their locomotion gait from swimming to crawling in response to changes in the viscoelasticity of the surrounding fluid.^24^ We found that the relative contributions of all but one of the eigenworms exhibit a clear increasing or decreasing trend with gel concentration, supporting a smooth behavioural gait modulation (Fig. 2g) and suggestive of the changing relative costs of the different modes in different environments.^35^ The low dimension of postural space reveals the highly constrained biomechanics of gaits and manoeuvres, available to *C. elegans* even in 3D and across a range of media.

**Figure 2:**
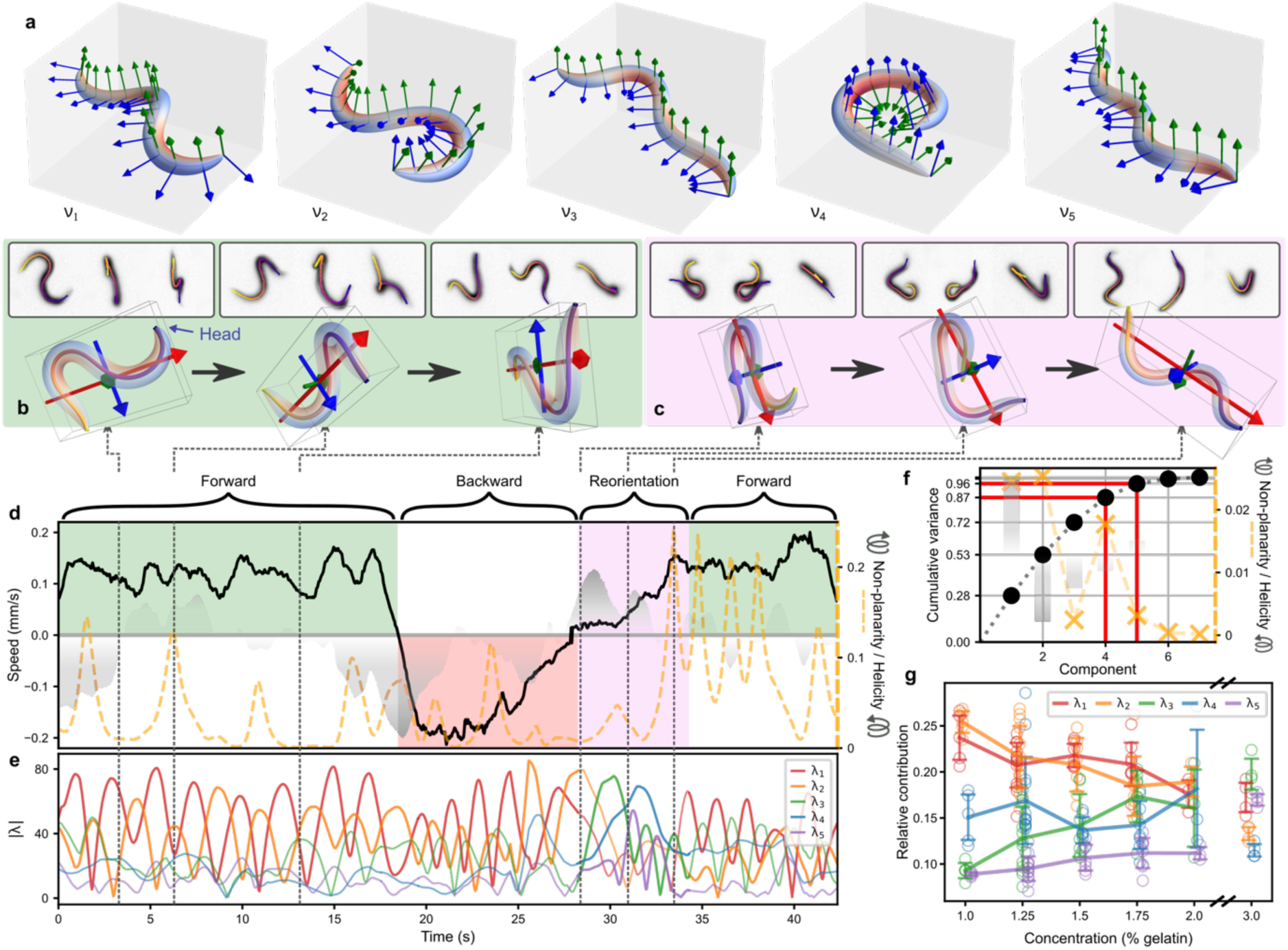
Non-planar postures give rise to non-planar locomotion. a. First five eigenworms (ν_1_-ν_5_) of 3D postures. Body colour indicates the signed curvature (Methods). b-d. A 40-sec clip (d, Supp. Video 3 Ch. 3.6) including forward locomotion, reversal and a manoeuvre, showing sample frames (three views) and body shapes (b-c). The three principal axes of the body (red, blue and green arrows) form the encompassing boxes. b. Forward locomotion alternates between near-planar and helical postures to reorient the undulatory plane without slowing. c. A turn begins with an Ω-body-shape (left) but ends with the head projecting out of the body plane (centre) to resume forward locomotion along a new trajectory with a helical posture (right). d. Kinematics reveal spikes in posture non-planarity (orange dashed line) and alternating helicity of postures (grey shading) that do not compromise speed (black line). e. Eigenworm coefficients (λ_1_-λ_5_). λ_1_ and λ_2_ oscillate during forward and backward locomotion. Contributions from higher-order modes peak during turn manoeuvres. f. Five eigenworms account for 96% of the variance in postures across all conditions (black spots). ν_1_ and ν_2_ are highly non-planar (orange) and helical (shading), with opposite handedness. g. Relative eigenworm contributions in fluids with increasing resistivity: λ_1_, λ_2_ dominate coiling gaits, characterised by shallow waveforms; λ_3_ and λ_5_ increase during 3D crawling, characterised by high-curvature short-wavelength undulations; λ_4,_ most strongly associated with manoeuvres, shows no apparent concentration dependence. Mean: solid lines; single trials: open circles; error bars: ±1 std. P-values (Pearson correlation): λ_1_-λ_5_: 2.99×10^-4^, 2.20×10^-7^, 1.45×10^-5^, 8.16×10^-2^, 2.23×10^-12^.

Similar to planar undulations, in 3D *C. elegans* propagates bending waves along its body (Ext. Fig. 1, Supp. Video 2). However, the resulting postures can be highly non-planar (Methods, Fig. 2b-d). We identify distinct locomotion gaits (Supp. Video 2): *clockwise coiling, counterclockwise coiling,* an *infinity* gait (collectively: *chiral* gaits) and 3D crawling.^36^ 3D crawling combines planar undulations with frequent twisting of the body into 3D shapes, often helical, and rotation of the body in 3D, which we call rolling (Supp. Video 2 Ch. 2).

Coiling gaits employ sequences of nearly planar postures with low helicity (Methods), in which the head moves in a clockwise or counterclockwise elliptical trajectory (Supp. Video 2 Ch. 3).^36^ Infinity gaits involve alternating clockwise and counterclockwise phases of the head and neck motion in each undulation.^36^ While chiral gaits are more prominent in low-viscosity fluids and 3D crawling at high viscosity, all gaits are observed in all environments. Furthermore, while animals can maintain gaits for extended durations, they frequently transition between and even mix different gaits, both in forward and backward locomotion (e.g. Supp. Video 1 Ch. 2.1). As all these crawling and 3D coiling gaits involve non-planar postures, we call them non-planar gaits.

Coiling (Supp. Video 2 Ch. 3) and 3D crawling (Supp. Video 2 Ch. 2) are distinct behaviours. Coiling is a low-speed gait, characterised by high-frequency rolling of the entire body and straight-line motion of the centre of mass. The head and neck move elliptically with the body following.^36^ During 3D crawling, non-planar motion of the head and neck is more pronounced at the start of undulations (contrasting with continuous rolling during coiling gaits). As the crawling undulation wave propagates posteriorly, the body curls into helical postures (Fig. 2d) which can also smoothly steer the body out of the primary undulatory plane (Supp. Video 2 Ch. 2.2). Of course, perfect helices do not deform and hence cannot self-propel. In our data, despite the helical-looking body shapes, 3D crawling produces comparable speed to planar sinusoidal crawling (Fig. 2d), which can be explained by the high-amplitude curvature wave propagating along the animal (Ext. Fig. 1a). As animals move, they alternate between low-speed chiral gaits (Ext. Fig. 1b-d) and high-speed 3D crawling, both combining whole-body undulations with twisting initiated by the head and neck. Consistent with an intermittent (stop-and-go) search strategy, frequent pauses interrupt both locomotion and manoeuvres, but the head and neck often continue to scan the local environment (Fig. 3h, Ext. Fig. 4).

**Figure 3:**
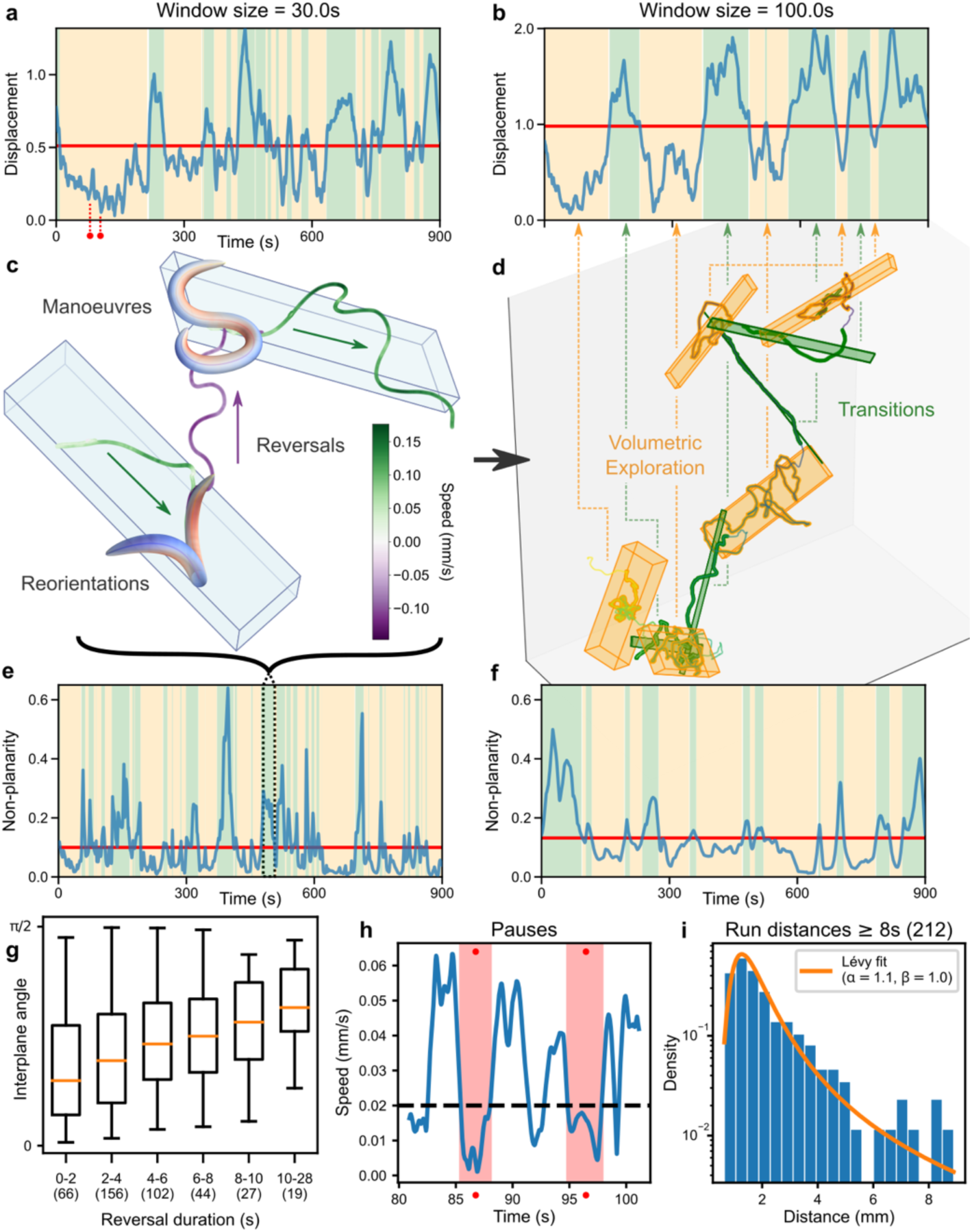
Locomotion primitives combine to produce a volumetric exploration strategy. a-b. Moving-average displacement reveals fluctuations across scales (mean, red line; >mean, green; <mean, yellow). c. Complex manoeuvres such as φ-turns result in a new local plane of exploration. d. Volumetric exploration consists of local patches (orange boxes) that include runs, simple turns and pauses, resulting in low displacement (b), and transitional, high-displacement runs (green boxes). e-f. Manoeuvres (complex or compound turns) produce non-planar trajectories with low displacement (a-b). g. Longer reversals produce higher inter-plane angles during subsequent φ-turns (n=414, total ∼1500 sec, making up 8% of total duration of the trajectories). h. Forward locomotion can be interrupted by brief pauses (red, see red time labels in a). We observe switching between higher-speed runs and lower-speed or stop-sense-and-go locomotion. i. Heavy-tailed run distance distribution across the dataset reflects many short local-search runs and occasional long (often transitional) runs.

We asked whether these non-planar locomotion gaits explain the observed volumetric trajectories (Fig. 1f-i, Supp. Video 1). We find that over tens of seconds, despite the non-planar postures, non-planar locomotion gaits and steering, animals often move in a shallow volume that we call quasi-planar patches (quantified by the non-planarity of the local trajectory, Methods, Figs. 2d, 3e). Local patch search has previously been associated with environmental features^7–11,37^ but as it arises here without any such constraints, it is perhaps more reminiscent of patch-like foraging strategies of mammalian swimmers.^38^ Why then, do *C. elegans* prefer quasi-planar patches, and how are long-term volumetric trajectories achieved? We conjectured that turn manoeuvres facilitate volumetric exploration. We found and labelled two categories of turning manoeuvres, *simple turns*, occurring during forward locomotion, and *complex turns*, preceded by a reversal (Supp. Video 3). Both simple and complex turns involve high-curvature body shapes, associated with increased contributions from the higher-order eigenworms (Fig. 2d,e). To characterise turn manoeuvres, we compute the principal axes of the local trajectory in a small window before and after each turn and define two turn angles: trajectory turn angles between segments of the trajectory and inter-plane angles between the corresponding principal planes (Methods, Ext. Fig. 5). Simple turns are characterised by small inter-plane angles despite arbitrarily large trajectory turn angles (keeping the animal within the local quasi-planar patch), but the occasional occurrence of multiple simple turns in close succession gives rise to significant 3D reorientation (Ext. Fig. 6, Supp. Video 3 Ch. 2). By contrast, while complex turns vary in their details, they invariably involve 3D postures and change the locomotion plane (85% of complex turns change the locomotion plane by >36 degrees, Ext. Fig. 5e). In 2D, omega^5^ or delta^34^ turns, which similarly follow reversals, are so called in recognition of the observed (Ω or δ-like) posture at the peak of the turn. In 3D, we label complex manoeuvres φ-turns, reminiscent of the worm’s posture as it exits the turn (Fig. 2c and Supp. Video 3 Ch. 3).

Turn manoeuvres allow worms to alter their heading more than steering alone (Fig. 3c). In particular, the stereotypic φ-turn sequence (forward-reversal-turn-forward) enables large changes to the trajectory direction both within the local plane of exploration, and between the local planes before and after the turn (Methods, Ext. Fig. 5).^39,40^ Longer reversals are correlated with larger inter-plane turn angles (Fig. 3g), suggesting that the time penalties incurred by reversals are necessary to explore the volume. Clusters of simple turns, which we dub compound turns, can change the local plane but incur even greater time penalties (Ext. Fig. 6).

On timescales of minutes, animals exhibit significant speed variability, including prolonged periods of low displacement (Fig. 3b) and high nonplanarity (Fig. 3f), characterised by clusters of turns and extended periods of resting accompanied by sensing (Fig. 3h, Ext. Fig. 4). Steering, simple turns and complex manoeuvres (Fig. 3c) enhance local search, limiting dispersal. Occasional long runs (i.e., periods of forward locomotion between turns) take animals from one local search region to new, unexplored ones (Fig. 3d). These long runs make up the tail of a Lévy distribution across all run durations (Figs. 3i,4b), reminiscent of Lévy-flight foraging strategies.^7,41,42^ Taken together, volumetric foraging trajectories emerge from combinations of 3D gaits, steering and simple turns which produce quasi-planar (far from isotropic) patches; these patches are separated by complex and compound turn manoeuvres and stretches of forward locomotion. The richness of behaviours on different timescales, and the many decision points during this exploratory behaviour suggest constant trade-offs between local sampling and exploration, translating into trade-offs between planar and non-planar local actions.

**Figure 4:**
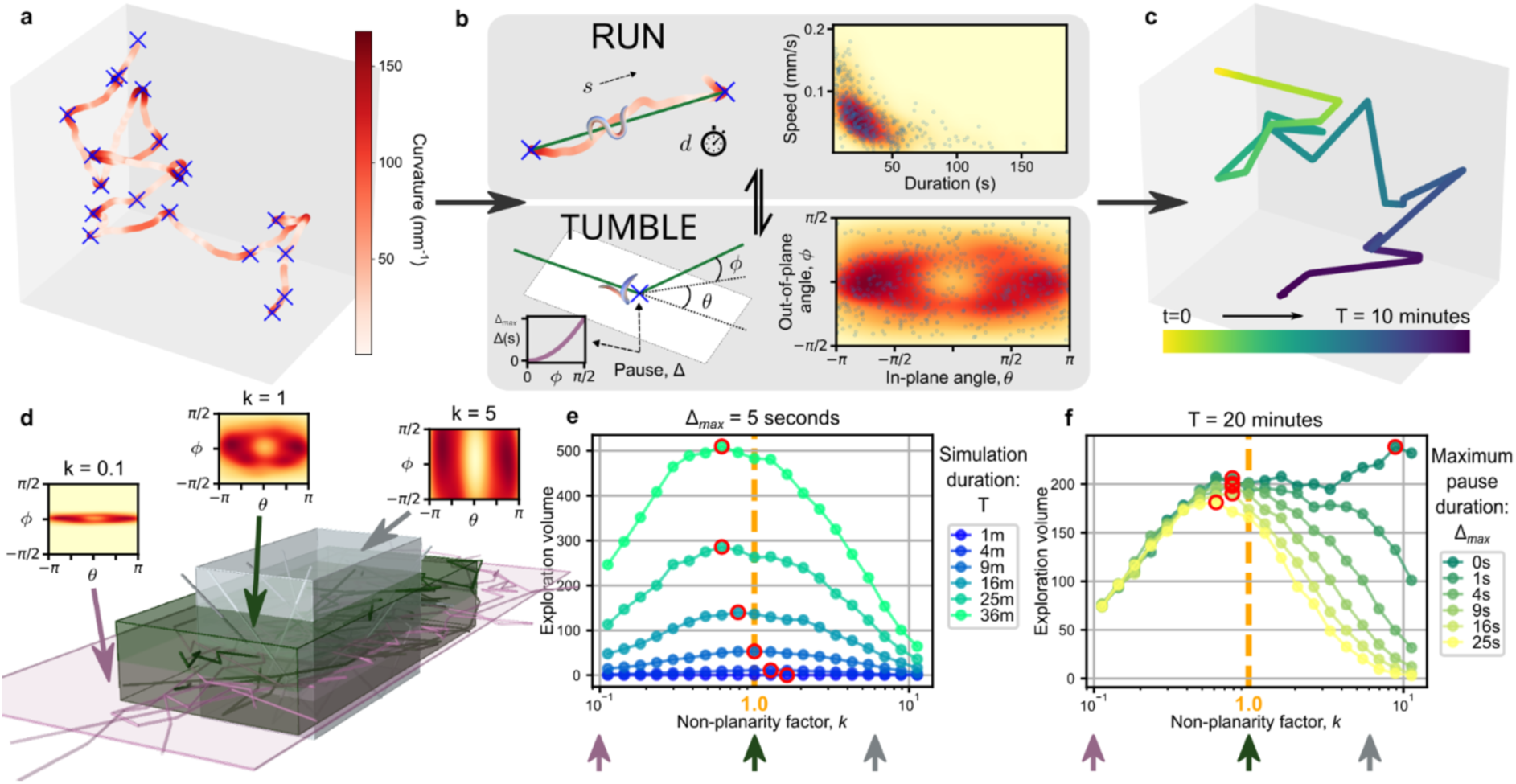
A quasi-planar strategy is optimal for *C. elegans* volumetric local search. a. A run-and-tumble approximation (blue crosses) superimposed on a 10-minute empirical centre-of-mass trajectory. b. Point trajectories are generated from straight-line segments (runs) and turns (tumbles). Run and tumble parameters are sampled from two joint probability distributions, constructed from empirical data (blue points, n=513 runs, n=565 tumbles; heatmaps generated from calculated pdfs). Simulated point animals sample a straight-line run, move along this run in time and space and then sample tumble parameters. The animal pauses at each tumble by a duration, proportional to ϕ (Methods). c. Example simulated trajectory. d. The volume explored is approximated with a minimal encompassing cuboid aligned to the PCA components of the trajectory (Methods). By varying the out-of-plane turn angle distribution (through the non-planarity factor, *k*), trajectories vary from being long and planar (e.g. for *k* = 0.1) to short and volumetric (e.g. *k* = 5). e. Optimal *k* values (red circles) approach empirical fit (orange dashed line, *k*=1) for relevant simulation durations. Average over 1,000 simulations for each data point. f. Optimal *k*-value depends on turn penalties.

### Worms use an optimal strategy to explore 3D space

In models of random walks, from Brownian motion to run-and-tumble motion, particles travel the same distance along their paths, whether their trajectories are planar or volumetric, but planar exploration increases the chance of returning to the same proximity. We therefore asked what cost-benefit considerations might account for *C. elegans*’ quasi-planar exploration strategy. In analogy with the principle of maximum entropy, in which systems maximise their entropy (or information gain) subject to energetic constraints, we conjectured that *C. elegans* choose actions to maximise their exploration volume. A key consequence of the maximum entropy principle is that different states or system configurations are sampled according to the Boltzmann distribution, with decreasing frequency for increasing cost. In all species, locomotor actions can be described by a hierarchy of costs. In many species, and in *C. elegans* in particular, we hypothesise that this hierarchy is best described by the dimensionality of the motion: 1D approximately straight-line motion, 2D quasi-planar motion, and 3D volumetric motion. We therefore posit that *C. elegans* optimises its foraging strategy by sampling actions with frequencies matching this hierarchy, in accordance with the principle of maximum entropy.

To statistically characterise the data, we first sought to establish whether turn penalties are consistent with the principle of maximum entropy. *C. elegans* uses steering and simple turns to explore quasi-planar patches of its environment but reserves costlier complex or compound turns for 3D exploration. Rather than energetic costs, we argue that *C. elegans* foraging is a time-limited behaviour: After a period of foraging, animals will abandon the search and disperse.^5^ Hence, during foraging, time may serve animals as the currency of energy. We found that the more volumetric the manoeuvre, the greater the time penalty (Fig. 3g). Furthermore, the longer the reversal, the rarer it is, with densities closely following the Boltzmann-distribution (Ext. Fig. 7). Next, we approximated empirical trajectories by straight-line runs and turning events (Fig. 4a, Methods). We find no evidence that the internal structure of runs (Fig. 3a,b,h) predicts the timing, type, or angles of its preceding or subsequent turn (Ext. Table 1), validating the two-state run-and-tumble approximation. We may therefore absorb variable-speed locomotion and pauses into a single run state and collapse simple and complex turns, including reversals, into a single tumble state. The dominance of runs over localised, infrequent turns points to a higher cost of turning over forward motion,^43^ heuristically consistent with the maximum entropy principle. The associated kinematic parameters reveal heavy-tailed run distance, duration and speed distributions (Figs. 3i, 4b), broadly distributed in-plane turn angles, θ, and narrowly distributed out-of-plane angles, ϕ, explaining the appearance of quasi-planar exploration punctuated by non-planar turns. Taken together, the foraging actions in 1D, 2D and 3D, and their statistics, suggest a compact strategy, underpinned by a compact decision-making circuit, explains the quasi-planar exploration strategy.

We next asked whether the statistics of runs and tumbles optimise the exploration volume. To test whether the statistics of the quasi-planar strategy used by *C. elegans* maximise exploration volume, we developed a parsimonious biologically-grounded model. We adopted a data-driven approach starting with the approximation trajectories and explicitly imposing the empirical frequencies and parameters of different actions (Fig. 4a-c, Methods). To construct model trajectories, runs are sampled from a joint empirical distribution of duration and speed, and tumbles from a joint distribution of in-plane and out-of-plane angles (Fig. 4b). To account for the correlation between the duration of the manoeuvre and out-of-plane turn angle, ϕ (Fig. 3g), at each tumble, the model worm pauses for a duration, Δ(ϕ) (Fig. 4b). The two free parameters of the model, the simulation time, *T*, and the maximum pause duration, Δ_max_, approximately match the empirical data. Combined, the data-driven approach ensures the implicit cost-frequency relation is preserved.

We asked what turn strategy maximises the explored volume (Fig. 4d-f). As non-planar turns incur time penalties in our model, a strategy that maximises exploration volume must balance between distance travelled and non-planarity of the trajectory (Fig. 4d, Supp. Video 4). Exploration strategies that produce the largest volumes closely match those of *C. elegans* (Fig. 4e,f, Ext. Fig. 8). The strategy is near optimal for simulation durations commensurate with typical local-search timescales (10-25 minutes^5^). More planar strategies fare better for sufficiently long times or if the delays associated with out-of-plane turns make them prohibitively expensive. In summary, *C. elegans’* hierarchy of costs for motion in one, two, and three dimensions results in a far-from-isotropic, quasi-planar exploration strategy, producing a near-optimal volume-coverage of 3D space, as predicted by the principle of maximum entropy.

## Discussion

We presented the first corpus of 3D animal locomotion and foraging in a homogeneous environment, revealing new non-planar gaits and manoeuvres. The φ-turn appears to generalise the familiar omega and delta turns used by *C. elegans* on a substrate. Several observations support this conjecture. Like φ-turns, omega and delta turns follow similar stereotyped sequences of forward locomotion, reversal, turning, and a resumption of forward locomotion. Omega and delta turns recur in contexts of search,^5,8^ navigation^2,44^ and escape,^45,46^ and are underpinned by a common neural circuit,^5^ suggesting that φ-turns too are similarly controlled. The reversals in all three incur a substantial time penalty to the animal. Despite the apparent similarity of forward and backward locomotion gaits, their detailed kinematics differ, and hence reversals likely incur a greater metabolic cost as well.^47^ Combined, the costs of reversals may dominate the cost of the high curvature posture during the turn.^48^ Reversals preceding omega and delta turns have previously been motivated purely by the need to rapidly avoid danger. We find that the φ-turn confers an additional benefit for volume exploration, with longer reversals leading to larger 3D turn angles, and conjecture that these manoeuvres have specialised for directed out-of-plane turning. Combined, our observations suggest that on a substrate, *C. elegans*’ attempted φ-turns are intended as full 3D manoeuvres but are frustrated by being physically constrained to the substrate, manifesting as Ω-or δ-shaped turns.

While nematodes typically do not inhabit homogeneous gelatin without sensory cues, our results present an important baseline for the study of 3D locomotion primitives and exploration strategies, which extends previous work, in which gravity and the physical structures in the environment impose biases on the exploration strategy and the animal’s neural representation of the volume.^37,38,49,50^ By simultaneously tracking postures and trajectories in 3D, we found a foraging strategy that highlights the trade-offs between local sampling versus moving, and exploration distance versus space or volume coverage. Our finding of trajectories comprising quasi-planar local patches generalises planar hierarchical and Lévy-flight exploration strategies^7–11^. The optimal volume covering strategy during *C. elegans* 3D exploration reveals more fundamental and universal principles of navigation and search that arise from the need to maximise information under time and energetic constraints. The emergent strategy then samples actions according to a hierarchy of costs associated with 1D, 2D and 3D embodied locomotion. Deciphering how neural circuits for decision making and motor control orchestrate the rich repertoire of *C. elegans* behaviours in 3D will elucidate the encoding of optimal decision-making strategies in simple brains.

## Methods

### Setup

We built a bespoke triaxial imaging system consisting of three approximately orthogonal cameras (Ximea xIQ MQ042RG-CM), each connected to a variable-magnification telecentric lens (minimum 7X, Navitar Inc.). Each camera has a pixel size of 5 μm and records 2048×2048 pixel resolution images. The cameras were synchronised using a trigger box. Videos were recorded using the StreamPix 6 software (Norpix) at 25 frames per second and concurrently written to separate hard drives.

The telecentric property of the lens means the light rays are close to parallel (±0.4° error), producing the large depth of focus. The lens magnification was set to ensure sufficient resolution with a depth of focus of at least 1cm and is thus suitable for capturing postural sequences and exploration trajectories over several minutes.

A glass, fluid-filled sample cube was backlit with three diffused red LED lights located opposite each camera (Fig. 1). Cubes varied in size (minimum 2 cm side length), with larger sizes used to ensure there are no observable wall effects.

### C. elegans growth and maintenance

Wild-type (N2 Bristol strain) *C. elegans* worms were grown at 20°C and maintained under standard conditions.^51^ Worms were age-synchronised^52^ and observed as young-adults.

### Experiments

1%-4% (v/v) gelatin solution in M9 buffer was prepared to the appropriate concentration as per Berri et al.,^24^ heated and loaded into the sample cube when molten. The cameras were then immediately calibrated, and the gel was then left to set and cool to room temperature (20°C) prior to behavioural video recordings. Concentrations of 0.5% and lower were excluded due to worms visibly sinking in the fluid. Calibration slides were prepared by printing an oriented grid pattern onto glass coverslips. A calibration slide was positioned in 100-200 randomly chosen orientations in the fluid-filled cube. For each slide position, a triplet of calibration images was captured simultaneously by the three cameras. Using OpenCV pinhole camera models,^53,54^ these images were used to determine the initial camera-model parameters.^36,55^

Each trial consists of an individual worm picked using a platinum wire and placed in the sample cube, approximately in the centre of the field of view. The recording is started after at least 30 seconds to let the worm settle and continued while the animal is visible in all three camera views. Clips that last at least 4 minutes were included in the corpus (numbering 52 clips, and with a maximum duration of 25.5 minutes). Since the sample cube was always larger than the volume of view, a small number of trials were sometimes conducted using the same experimental preparation and, consequently, there were occasionally more than one animal in the recordings, though typically only the latest animal deposited in the cube was in focus in all three views.

After 3D trajectory tracking from the recorded movies, we used the calibration images to impose world coordinates on the trajectories and verified that animals were not sinking in the gelatin due to gravity.

### Video preparation

Areas of interest (containing worms) were identified in each view with custom software (using OpenCV^53^) and the remainder of the images were replaced with a static background which was computed using the median pixel values across the clip. Lossless encoding (H264 codec) of the new image stack (“compressed videos”) was used to reduce terabyte-sized recordings to megabytes.

### 3D tracking

We developed segmentation and tracking software that can be applied either to compressed or raw videos. Candidate worms were identified in each image using thresholding and contouring functions in OpenCV^53^. For each contour in every frame, the 2D centre of mass was calculated. Calibrated pinhole camera models were used to triangulate objects (defined by these contours) across the three views and find corresponding 3D centre-of-mass coordinates using an iterative optimisation algorithm to minimise the reprojection error to all images (L-BFGS^56^). Specifically, objects were identified by contour combinations with error capped at 5 pixels between the 2D centre of mass and the reprojection of the 3D centre of mass onto the same images. Consistently projected objects trace out a tracking trajectory over time. In many cases, additional objects were present in one or more views (whether transient bubbles, other worms, tracks, etc.). Most of these were handled automatically, by discarding candidate points that are inconsistent with the trajectory up to that time. When the worm traverses an obstruction such as a bubble along its trajectory, that was not resolved automatically, the tracking trajectory was guided manually for up to a handful of frames. Where posture reconstructions are available (see *3D posture reconstruction and camera parameter adjustment* below), final trajectories use the centre-of-mass of the body points instead of the above procedure. Finally, trajectories are smoothed with a 1 second moving average.

### Intrinsic curve representation

We modelled the midline of the worm as a 3D curve using a Bishop frame parametrisation that naturally admits biologically-informed constraints and regularisation.^57^ The Bishop frame is independent of location and orientation in the laboratory frame coordinates. Previously, the Frenet frame has been used to represent *C. elegans* body shapes.^23^ However, to avoid numerical issues for points of low curvature (and ill-defined normal vectors when the curvature is zero), Stephens et al.^23^ represented the curve in terms of the bending angles, rather than the curvature.

Bishop^57^ proposed an alternative to the Frenet frame that is well-defined and well-behaved for all curvature values, albeit with a rotational degree of freedom (i.e. it involves an arbitrary choice of rotation of the frame around the tangent). Like the Frenet frame, the Bishop frame consists of three unit-vector fields (the tangent *T* and two normals, *M*_1_ and *M*_2_), forming an orthonormal moving frame along the curve:

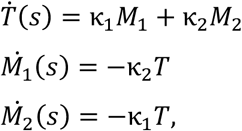

where the dot denotes differentiation with respect to the arclength *s* and κ_1_, κ_2_ are the two components of the curvature in the *M*_1_ and *M*_2_ directions, respectively (Ext. Fig. 2a,b).

The generalised vector curvature 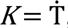, pointing perpendicular to *T* at every point along the curve (*s*), can then be represented as the sum of curvature components κ_1_ and κ_2_ along the frame vectors *M*_1_ and *M*_2_, respectively: *K* = κ_1_ *M*_1_ + κ_2_ *M*_2_. The curvature magnitude, κ ≡ |*K*|, defined at every point along the curve, is agnostic to the choice of frame, and can hence be obtained directly from the reconstructed postures. When visualising the curve, we chose a primary curvature direction as positive, thus obtaining a signed curvature from the vector curvature *K*.

### 3D postural reconstruction and camera parameter adjustments

As a worm moves, the image focus changes between the three views, making edge or feature matching impractical. To best capture body shapes as viewed by each camera, posture reconstruction software was developed to jointly optimise camera parameters of our Pinhole camera models alongside the curve parameters representing the worm midline. Rather than feature or edge detection, we developed a differentiable renderer to construct the full-body images from 2D projections of the 3D curve. The rendered images were compared with the recorded images to generate a pixel-wise error. The combined loss was minimised using gradient descent (Adam optimiser^58,59^) with a decaying learning rate until convergence criteria have been met.

Only the first few frames were used to update our pinhole camera models from an initial model constructed from the calibration images. For the rest of the video, continuous adjustments during the optimisation were limited to the two offsets in each camera model (associated with its two coordinate axes) to allow more precise reconstruction. Minor adjustments of these offsets accounted for small optical variations over time, while the main camera model parameters are fixed to prevent parameter drift in the reconstructions. Subsequent frames were instantiated with the solution from the previous frame for efficiency and consistency.

Many of the tightly coiled postures cannot be estimated accurately, even manually. As the worm returns to simpler shapes, several interim frames may be incorrectly reconstructed and even the head-tail orientation reversed. Following the initial pass, head-tail flips were manually identified and corrected. Then incorrect frames or short sequences were patched using an adjusted loss that blends the main loss with a continuity cost for all parameters to ensure smoothness at the stitch points. The method was benchmarked against other 3D reconstruction algorithms.^36,55^ For more details see Ilett et al. (2023).^54^

### 3D eigenworms

Principal Component Analysis decomposition of planar worm postures is well established,^23^resulting in a four-dimensional basis of eigenworms that describes the repertoire of *C. elegans* locomotion on agar. We extended the eigenworm formulation into the complex domain such that the resulting basis consists of 3D eigenworms; the method is known as complex PCA^60,61^ (denoted here cPCA). cPCA defines a unique representation of a curve (here, the body shape) that is independent of location and orientation, and hence agnostic to the anatomical (lateral and dorsoventral) information, which we do not capture. Using cPCA, we constructed a low-dimensional shape-space embedding of 3D postures observed across all gel concentrations in our dataset.

We start with a curve of fixed length represented by the curvature components (κ_1_, κ_2_), obtained in the corresponding Bishop frame (see above). In the complex domain, we may represent each body shape as *w* = *w*(*s*) as w(*s*) = κ_1_(*s*) + *i* κ_2_(*s*). Using this representation the body shape covariance is also complex and is given by

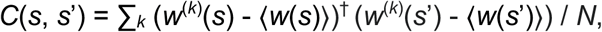

where *w*^(k)^ is the *k*^th^ body shape from a corpus of size *N*, ⟨·⟩ denotes the mean over all postures in all animals and ^†^ denotes complex conjugation. We wish to represent the body postures using our midline reconstructions (with 128 equally spaced points). After this discretisation, the covariance becomes a square matrix. The eigendecomposition of the covariance matrix, *C,* then yields an orthonormal basis of 128 eigenvectors (here called complex eigenworms) ν*_i_* that is unique up to arbitrary rotations of any of the basis vectors. In this basis, a worm’s body shape is given by a superposition of eigenvectors *w* = ⟨*w*⟩ + ∑*_i_* λ*_i_* ν*_i_* where the coefficients λ*_i_* = (*w -* ⟨*w*⟩) ν_i†_ include both amplitudes and relative rotations of the different components (Ext. Fig. 2c-e).

Due to the rotational freedom in the Bishop frame,.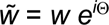 produces the same underlying curve as *w* for any rotation of the *M*_1_ and *M*_2_ frame around the tangent *T* by Θ (Ext. Fig. 2c). A consistent choice of the free rotation (also called phase normalisation) is required for the mean posture to be meaningful.^60,61^ Here, we chose the free rotation consistently so that the *M*_1_ direction captures the primary curvature direction. This choice does not explicitly account for the anatomical orientation of the animals (inaccessible in our data). To ensure that our choice does not bias the solution for our data, we tested an alternative approach, by performing off-centre PCA, in which the covariance is replaced by 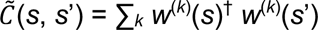. For our dataset, any normalisation or using off-centre PCA gives almost identical numerical solutions, as the mean posture is almost zero and rotations of the body shapes do not change 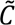 (since *e^-i^*^Θ^ *e^i^*^Θ^ = 1).

To assess gait modulation, a linear regression was performed on mean coefficients of each eigenworm as a function of the gel concentration. In this paper, we only consider the first five eigenworms which account for 96% of the variance across our corpus.

### Non-planarity

We define a non-planarity metric of postures and trajectories in terms of their PCA decomposition in the lab frame. First, we compute the three principal components obtained through PCA decomposition of either body or trajectory coordinates in the lab frame. For *n* > 2 points along the body for posture, or along a point trajectory over a time window, PCA produces an orthonormal basis (*n*_1_, *n*_2_, *n*_3_) and singular values *s*_1_ ≥ *s*_2_ ≥ *s*_3_. We define the normalisation factor σ^2^ = ∑_i_ *s*_i_^2^ / (*n*-1) such that the percentage of the variance explained by each of the components is: *r*_i_ = *s*_i_^2^ / ((*n*-1) σ^2^). We define our non-planarity metric as the ratio of normalised variances. 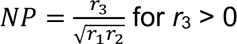 and 0 otherwise. The metric is valid for all curves, bounded below by 0 for planar curves (*s*_3_ = *r*_3_ = 0) and straight lines (where also *s*_2_ = *r*_2_ = 0) and bounded above by 1 (*s*_1_ = *s*_2_ = *s*_3_ and *r*_1_ = *r*_2_ = *r*_3_). It is scale and rotation invariant and hence can be used to compare between both postures and trajectories of varying lengths.

### Helicity

A helix is defined as a smooth curve with constant curvature and torsion. As curvature and torsion increase, the radius and pitch of the helix decrease to form a narrow, tightly wound coil. We define a notion of helicity, ζ, based on the product of the torsion with the curvature squared.^62^ To ensure our helicity metric is well defined for all curves, we express the result in terms of the curvature components in the Bishop frame:

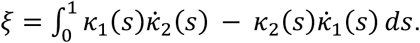

When considering the helicity of body postures, we normalise body lengths to 1 before the calculation. We choose the convention in which a positive helicity corresponds to right-handed chiral curves.

### Turn angle quantification for simple and complex turns

Turn analysis is performed for all simple and complex turns. Simple turns were obtained from the approximation trajectories (see below). We identify complex turns by reversals traversing at least 0.1 mm and establish 3-second incoming and outgoing trajectory sections on either side of the reversal (Extended Data Figure 5). Simple turns are similarly decomposed with two 3-second trajectory sections: incoming and outgoing. In addition, we identify compound turns (clusters of simple turns): two or more simple turns that occur within 5s (or within 10s, 15s and 20s for clusters of at least 3, 4 and 5 turns, Extended Data Figure 6). We characterise turn angles for each cluster by the incoming and outgoing trajectory sections to the overall cluster. This analysis is limited to the 43 experiments for which full posture reconstruction is available (yielding statistics for 1276 simple and 192 complex turns).

For each incoming or outgoing trajectory section, we first define a local reference frame in terms of the three PCA components of the local trajectory (Ext. Fig. 5). We then define two turn angles for each turn. The first principal component points in the direction of motion, and together the first and second components define the primary plane of the local trajectory.

Thus, we define the trajectory turn angle as the angle between the first principal components of the incoming and outgoing sections of the turn (or cluster). Angles between the third components of the incoming and outgoing sections (normal to the planes formed by the first two principal components) define the inter-plane (IP) angles.

### Speed and pauses

Worm velocity is defined as the derivative of displacement along the 3D trajectory, computed numerically. The speed is the magnitude of the derivative with respect to time. When full posture data is available, negative speed can be identified by comparing the velocity with the time-averaged head-tail alignment. Different types of pauses interrupt the locomotion. For a given behaviour or gait, speed informs the associated benefit and cost. However, *C. elegans* use different forward locomotion gaits, associated with different costs, frequencies, and speeds. Choosing a slower (chiral) gait might indicate that such gaits are less costly per undulation (as suggested by the smaller extent of the deformation along the body Ext. Fig. 1), or that such gaits confer additional benefits, e.g. to sampling the local environment. Pauses have well-studied benefits in a variety of animals and contexts (e.g. avoiding detection^1^ and improved swarming^63^). In the context of foraging, pauses have been linked to metabolic constraints as well as enhanced sampling.^1,42,64^ In our data, pauses include very brief alternating forward and backward locomotion, sudden cessation of motion across the entire body prior to continued locomotion, coiling up of the posture, or the cessation of motion of the mid-body while the head or tail continue to explore. To capture this wide variety of manifestations, pauses are defined as periods where the centre-of-mass speed stays below 0.02 mm/s for at least 2 seconds (Fig. 3h).^39,65^ During many pauses, animals are still active, particularly through continued movement of the head and neck (Ext. Fig. 4). As both speed and activity occur on a continuum, these thresholds were chosen to offer an intuitive description of the observed behaviour.

### Run-and-tumble approximation

Approximation trajectories convert empirical trajectories (see above) into run-and-tumble sequences of straight-line segments (runs) and point-turns (tumbles). In so doing, we collapse complex, compound and simple turns from spatially extended trajectories into a single point action, termed tumble. To remove any bias or artefact due to the variable speed along the trajectory, we begin by resampling the empirical trajectory to impose a constant speed. To divide the trajectory into runs and tumbles, we fix the start and end points of the trajectory and iteratively add candidate tumble points, which we call vertices, to the approximation. The approximation error is calculated as the mean squared distance in 3D space between every point along the constant-speed trajectory and every point on the approximation trajectory.

For each segment, we identify the candidate vertex that minimises the approximation error for that segment. We then select the lowest-error candidate vertex along the entire trajectory. Further vertices are added until the approximation error, calculated globally, falls below half of our error target (0.025 mm). Next, the vertices are pruned one at a time, with the least impact on the approximation error, until the error target (0.05 mm – a worm’s width) is reached. This process of adding and pruning vertices is iterated until convergence (i.e. the approximation vertices are stable). Finally, the vertices are mapped back to the corresponding points in the original variable-speed trajectory, and the corresponding run-and-tumble trajectory is saved. Run distances, durations and speeds are calculated directly from the straight-line segments between vertices (excluding end points). We fit the distribution of run distances by a Lévy distribution using the Levy-stable function in SciPy.^66^

To quantify turning angles at tumbles, we first calculate local planes for each incoming run, i.e., at each (non-endpoint) vertex, *i*, using vertices *i*-1, *i* and *i* +1. For a tumble at vertex *i* +1, in-plane (θ) turn angles are then calculated by projecting incoming and outgoing runs of vertex *i* +1 to the local plane; out-of-plane turn angles (ϕ) are similarly calculated for the projections out of the local plane (Fig. 4b). As these turn angles are derived from the approximation trajectories, they are mesoscopic, and hence differ subtly from the trajectory- and inter-plane turn angles defined above for simple and complex turns: both θ and ϕ are calculated at vertices between runs, rather than for local trajectory sections around the turn; and both θ and ϕ are calculated relative to planes that are defined non-locally (based on a number of vertices) rather than using the PCA decomposition of the local incoming and outgoing trajectory segments around the turn. Because θ and ϕ are defined purely from a run-and-tumble sequence of vertices, they serve as a useful representation, both for modelling and for comparisons with other datasets.

We use the approximation trajectories to identify simple turns, as any tumble excluding complex turns, and compound turns that are uninterrupted by at least one complex turn or reversal.

### Run-and-tumble explorer model

Model animals are represented as point particles that move along straight line runs and change direction through tumbles. Runs (defined by their speed and duration) and tumbles (defined by their in-plane and out-of-plane angles, θ and ϕ) are independently sampled using the parameter value distributions of the run-and-tumble approximations to the empirical trajectories. Each tumble consists of a reorientation coupled with a pause of duration Δ (in seconds), 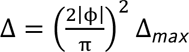 representing a time penalty incurred by the change in locomotion direction out of the local plane. The quadratic approximation effectively combines the negligible time penalty for simple turns and brief reversals preceding near-planar turns. When experimenting with the effects of non-planarity on exploration volume, we scale the ϕ distribution to smoothly modulate the strategy from planar to volumetric.

We wish to capture the empirical frequency of different actions, reflecting the underlying costs, as well as their key features, that determine the extent and direction of displacement over time. Run and tumble parameters are sampled from the data as follows. For each run, the speed and duration are jointly sampled from a bivariate Gaussian distribution fit to the empirical data. For each tumble, the tumble angles are sampled from the bivariate distribution of θ and ϕ. However, by definition, all turns have nonzero turn angles, the resulting bivariate distribution is annulus (or ring) shaped. Hence, we use a polar coordinate representation of the angles (*r*,α), where *r* reflects the overall magnitude of the turn angle and α captures the relative non-planarity of the turn, as follows. Assuming that the turn angle distribution is symmetric, we map the (θ,ϕ) values to their positive values. We split the α range [0, π/2] into 10 wedges and for each calculate a cumulative distribution function (cdf) of the *r* values. We also calculate a cdf for all α values. When sampling angles from this model, we first sample an α value from the global α cdf. Next, *r* values are sampled from the cdf of the corresponding α wedge and its neighbours. The final *r* value is a linear interpolation between the *r* values based on the α position within the wedge. The sampled (θ,ϕ) angles are computed by converting back to the Cartesian space and randomly applying a negative flip to each component.

We ran our model by generating simulated run-and-tumble trajectories. Each simulated trajectory starts with a worm at the origin facing a random orientation described by an orthonormal basis (*e*_0_, *e*_1_, *e*_2_). Runs and tumbles are then alternately sampled. First, a run is sampled and a straight-line is drawn in the *e*_0_ direction to a computed end point. Turn angles (θ,ϕ) are then sampled and the orientation is adjusted by a rotation of θ about *e*_2_ and ϕ about *e*_1_ to form the new orthonormal basis, (*e*_0_’, *e*_1_’, *e*_2_’). To estimate the 3D coverage of a trajectory, we define an exploration volume, *V* = *r*_1_ *r*_2_ *r*_3_, where *r*_1_, *r*_2_ and *r*_3_ are the maximum distances travelled along each of the PCA axes such that the exploration volume is a bounding cuboid aligned with the trajectory. Volumes are estimated by averaging over 1,000 simulations per data point (Fig. 4e-f, for standard deviations and standard error of the means, see Ext. Fig. 8). The volume estimate gives the same qualitative results as counting the numbers of voxels visited, with similar optimal values of *k* (Ext. Fig. 8).

To explore the turn strategies and their consequences for the geometry of search, we modulate the model worm’s exploration strategy with a non-planarity factor, *k*. Varying *k* modulates the out-of-plane turn angle distribution making trajectories more planar (ϕ→0) or more volumetric (uniformly distributed ϕ). The default, *k*=1, is obtained by fitting the empirical data (Fig. 4d). Parameter sweeps are performed over the two free parameters of our model: the maximum time penalty Δ_max_, and the simulation duration *T*, with sweep values slightly extending the range of the empirically observed values. In our data, complex turns are associated with reversals of up to 28 sec duration (Fig. 3g), and typical local search durations vary from minutes to tens of minutes (4-25 minutes in our data). Turns with negligible reversal durations, 0 < Δ < 2 sec, fail to produce volumetric exploration (with a lower mean inter-plane angle than a single simple turn, Fig. 3g). Small reversal durations, 2 < Δ < 4 sec, give rise to similar inter-plane turn angles as simple turns, with longer reversals (Δ > 4 sec) producing increasingly volumetric turns (Ext. Fig. 6). Sweeping over Δ_max_ tests the existence and robustness of the optimal strategy to different time penalties within the empirically observed range.

## Data availability

The data and analysis are available on https://doi.org/10.5281/zenodo.15460160. The repository includes all movies (after compression) and camera models, empirical trajectories, midline reconstructions, complex eigenworm decomposition of postures, kinematic metrics for postures and for trajectories, and sample scripts for visualizing the data. Included are: 52 movies (50 hermaphrodites and 2 male) with trajectory reconstructions (4-25 minutes per trajectory, totalling 6 hours 48 minutes). 3D midline reconstructions were obtained for 43 of the 52 recordings (4 hours 36 minutes).

## Code availability

Python software and scripts for data analysis, modelling and visualization are available on https://github.com/UoL-wormlab/wormlab3d. Sample scripts for loading and visualization of the data, as well as cPCA, are also available on https://doi.org/10.5281/zenodo.15460160.

## Acknowledgements

This research was funded by the Engineering and Physical Sciences Research Council (EP/J004057/1 and EP/S01540X/1), the Leverhulme Trust (ECF-2017-591) and a Leeds University Scholarship. We thank the University of Leeds for the use of the High-Performance Computing facilities.

## Author contributions

Conceptualization: NC, TPI and TR; methodology: DCH, FS, NC, OY, RIH, TPI and TR; software: FS, TPI and TR; formal analysis: TPI; investigation: FS, OY, RIH and TPI; writing – original draft: TPI; writing – review & editing: NC, TPI and TR; funding acquisition: NC, DCH, TPI and TR.

## Extended Data Figures

**Extended Data Figure 1:**
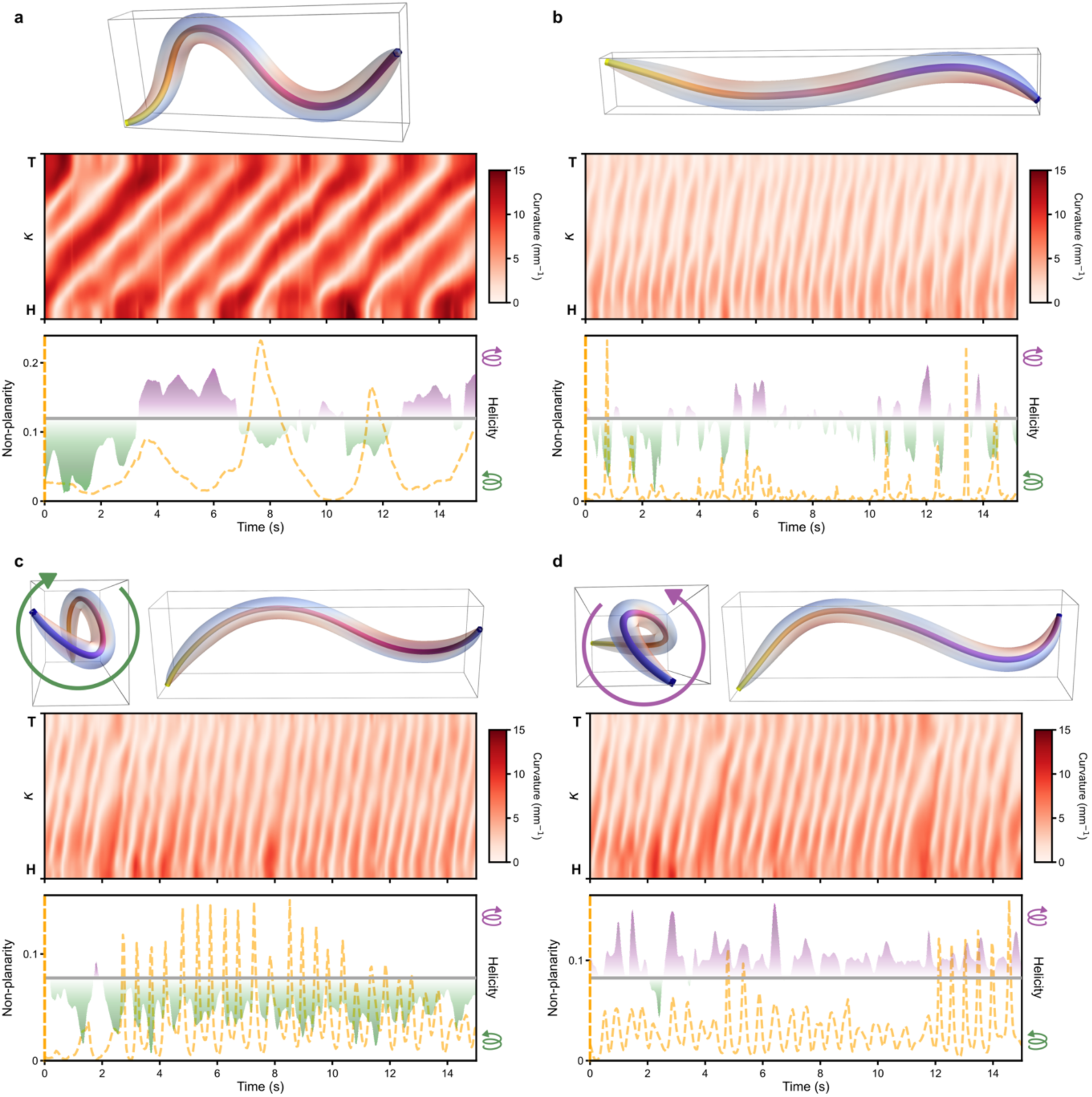
Kymograms for different 3D forward locomotion gaits. a. As in 2D, forward crawling in 3D consists of retrograde waves of high curvature magnitude propagating from the head to tail. Examples of chiral gaits including: b. infinity, c. clockwise coiling, and d. counterclockwise coiling. During coiling (c-d), low-curvature non-planar undulations propagate opposite the direction of motion. Characteristic frontal and sideways postures accompany kymograms, helicity traces and nonplanarity traces.

**Extended Data Figure 2:**
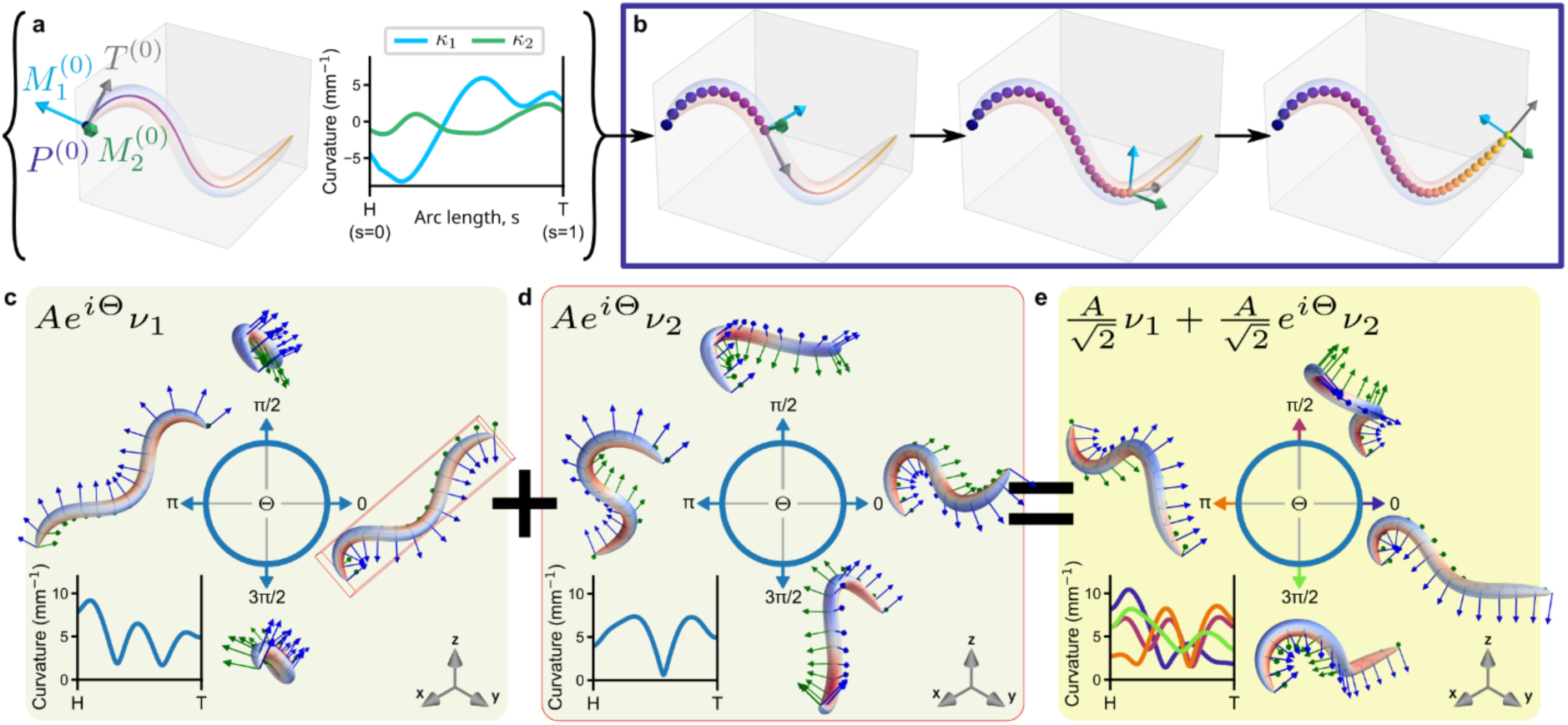
Decomposition of 3D postures into complex eigenworms. a. Reconstructed *C. elegans* body midlines are represented by 3D curves in the Bishop frame. The vector curvature of the curve, *K*(*s*) = κ_1_(*s*) *M*_1_(*s*) + κ_2_(*s*) *M*_2_(*s*), is decomposed into two orthogonal direction vectors, perpendicular to the tangent vector of the body midline, *T*(*s*). The curvatures, κ_1_ and κ_2_ are sufficient to describe a curve up to a translation and rotation in space. Without loss of generality, we impose a Bishop Frame (*T*^(0)^, *M*_1_^(0)^, *M*_2_^(0)^) on one end of the curve (*s* = 0), which we denote an initial point, *P*^(0)^. b. The body midline is constructed by integration of the Bishop equations from the initial point.^57^ c-d. Phase changes of a single eigenworm are equivalent to a rotation of the curvature frame around the curve, without changing its shape as shown in the insets. e. The linear combination of complex-valued eigenworms represents the worm‘s posture uniquely up to a global rotation. Generalising 2D (real-valued) eigenworms, in 3D, the weights (magnitude of the coefficients) and relative phases jointly determine each unique posture.

**Extended Data Figure 3:**
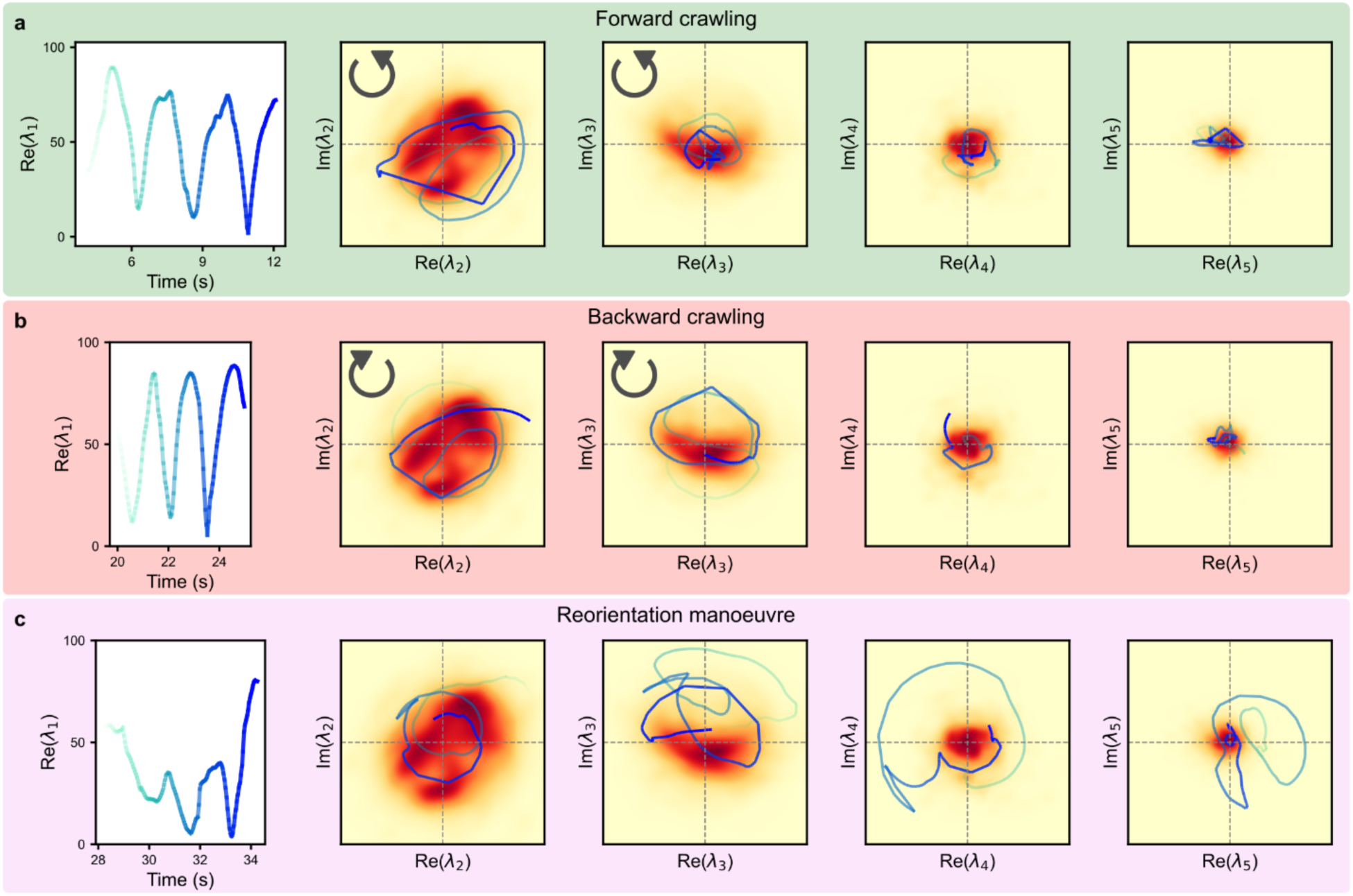
Clockwise and counterclockwise progression of the relative phases between eigenworms correspond to forward and backward locomotion, respectively. Without loss of generality the first eigenworm is rotated at each time point so that its coefficients lie in the real plane (left panels of a-c). Phase progressions (green to blue) of the coefficients of the complex eigenworms relative to the first coefficient (λ_1_), superimposed on corresponding density maps. The phases of λ_2_ and λ_3_ proceed counterclockwise relative to λ_1_ during forward locomotion (a) and clockwise during backward locomotion (b). The higher components, λ_4_ and λ_5_, play a lesser role during forward/backward locomotion (small distance from the origin) but grow during turn manoeuvres (c). The data shown correspond to the recording as Fig. 2d. Panels a-c match the first forwards (0-17.4 seconds), backwards (19.4-28.0 seconds) and reorientation manoeuvre (28.0-34.0 seconds) time windows in Fig. 2d with matching background colours. The density maps are the same across a-c and were computed from the coefficient values across the entire movie using a Gaussian kernel density estimation algorithm.^31,66,67^

**Extended Data Figure 4:**
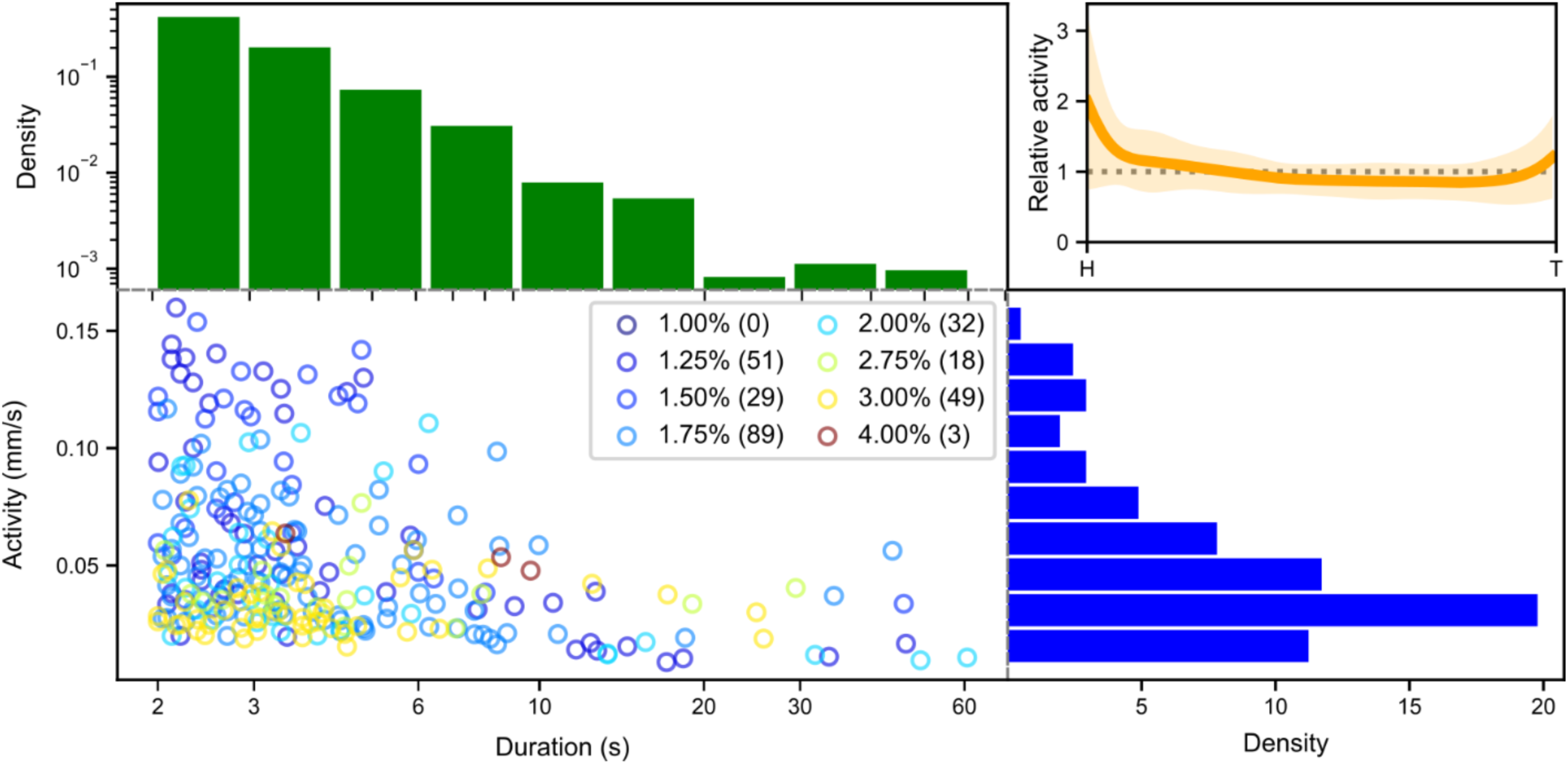
Pause durations and activity levels. Pauses can be brief (a few seconds) or extend to about a minute in our data, and their duration follows a heavy tailed distribution (green histogram). Pauses are observed across all gel concentrations apart from 1% with only 3 pauses in 4% (scatter plot). Elevated activity during a pause (blue histogram, defined as the mean speed of all body points, averaged over the duration of the pause), is due to parts of the worm continuing to move while the centre of mass remains stationary (<0.02mm/sec). This ongoing activity is mostly found in the head and neck (upper-right, mean activity ±2 std, relative to the whole-body mean). In 2D, head and neck activity are typically attributed to ongoing exploratory movements as the worm senses the environment.^68,69^

**Extended Data Figure 5.**
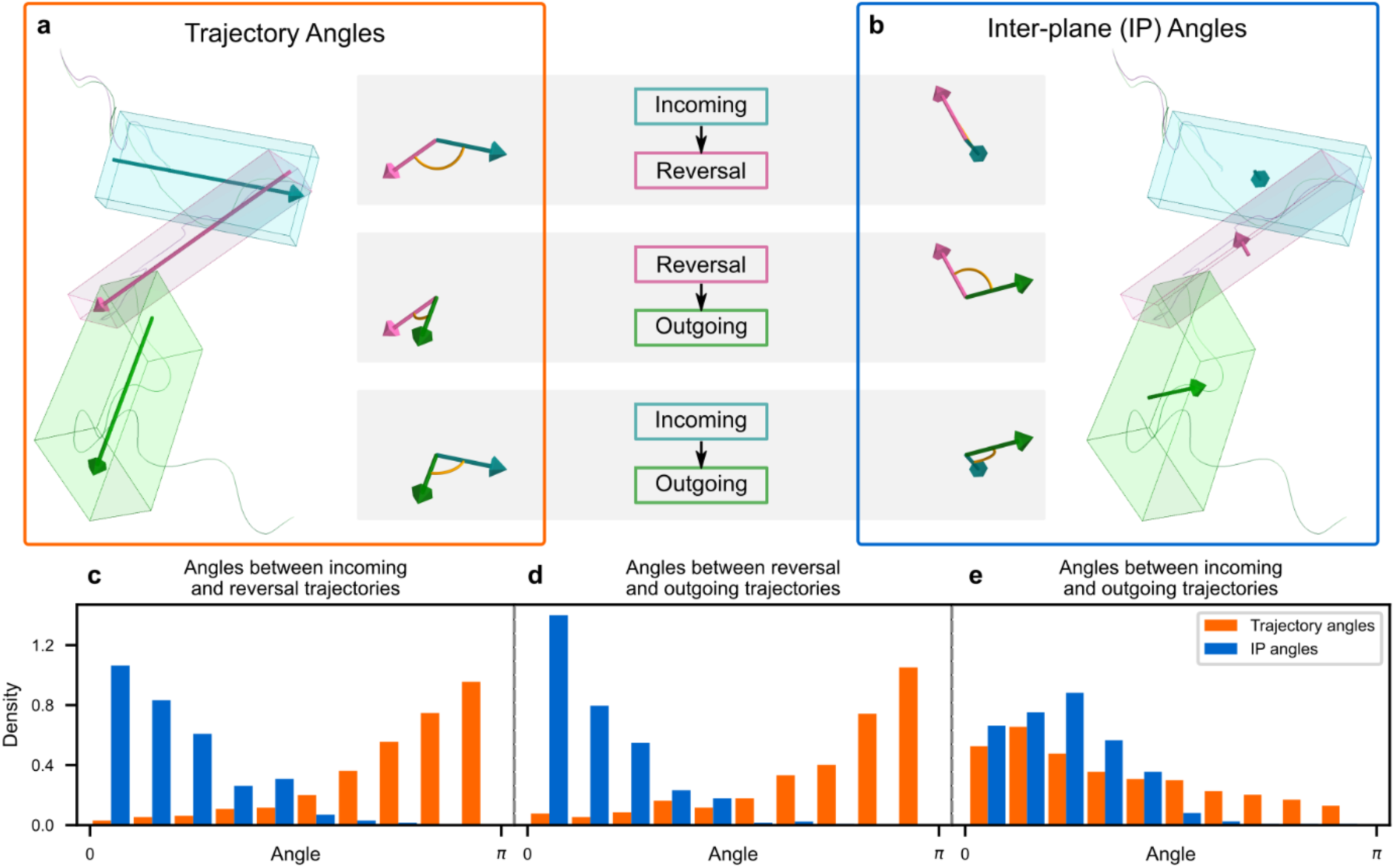
Three stages of a complex turn are required to generate 3D manoeuvres. a-b. We divide each complex turn into three sections: forward locomotion incoming (3s long, blue), reversal/backward locomotion (purple) and forward locomotion outgoing (3s long, green) and calculate trajectory turn angles (a, orange box) and inter-plane angles (b, blue box) between the sections. c-d. Analysis of all complex turns across our dataset (n=398) reveals that neighbouring sections are likely to have closely aligned axes with trajectory angles heavily skewed towards π and inter-plane (IP) angles heavily skewed towards 0. e. Both trajectory and inter-plane turn angles between the incoming and outgoing sections are more uniformly distributed, indicating that the reversal is as important as the transition from reversal to the outgoing trajectory for generating a new 3D direction.

**Extended Data Figure 6:**
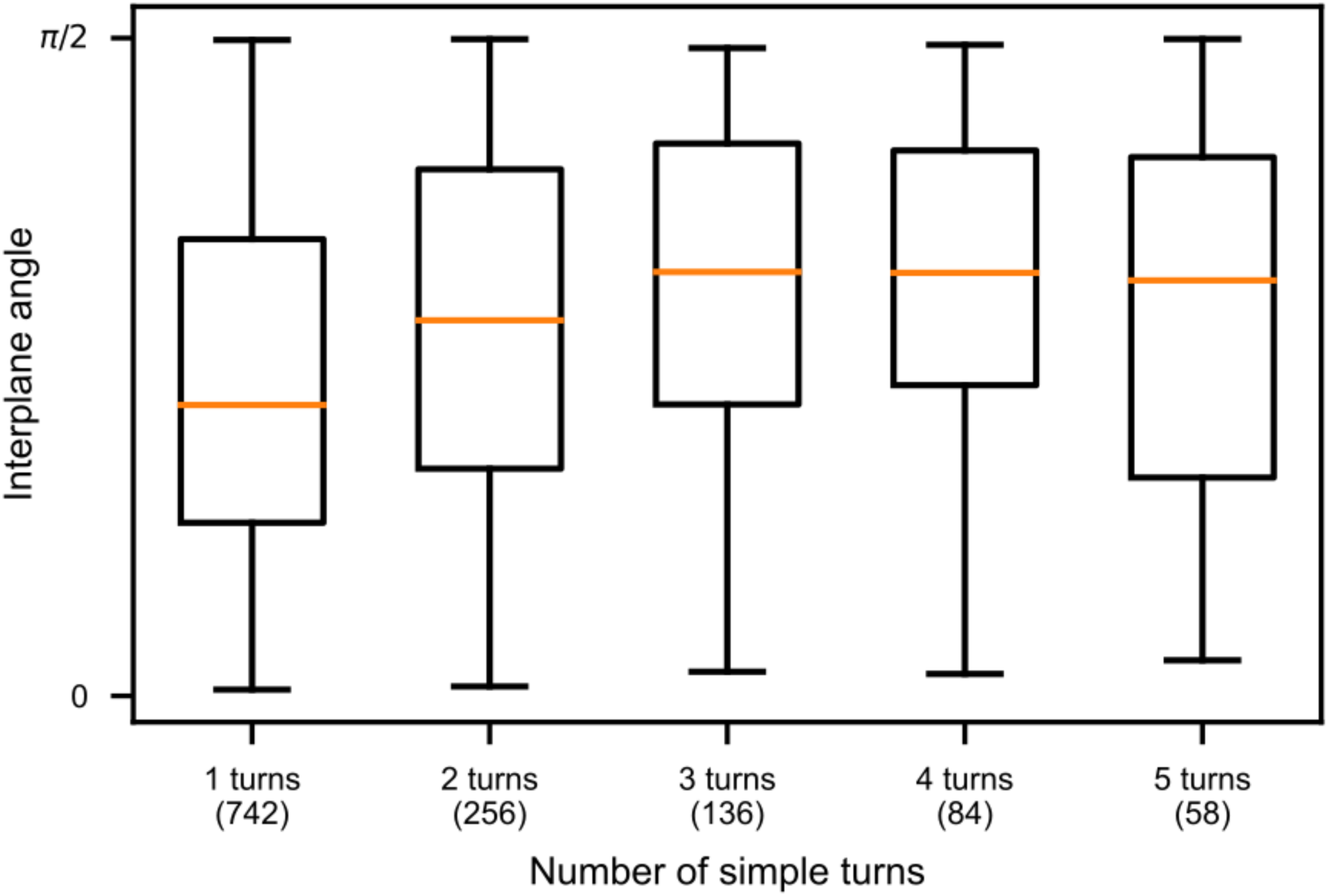
Compound turns induce large changes in the exploration planes. Compound turns produce 3D reorientation in a similar manner to a single complex turn (Ext. Fig. 5e). Two and three simple turns increase the mean inter-plane angle, akin to longer reversals preceding a complex turn. Event counts in brackets.

**Extended Data Figure 7:**
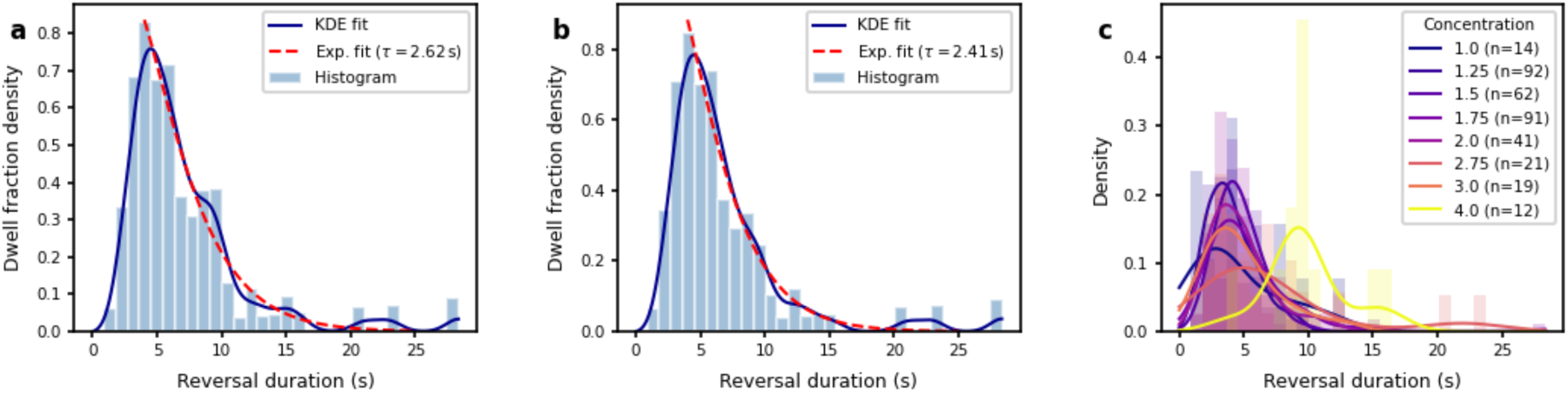
Reversal durations are well explained by a Maxwell-Boltzmann distribution. a. Distribution of dwell time density of reversals (reversal duration × normalised count) across all fully reconstructed movies (*n* = 352 reversals). An exponential fit (from 4-sec reversal duration, Exp. Fit) is superimposed on the histogram. Using the exponential fit as a null hypothesis and applying a Kolmogorov-Smirnov test, the null hypothesis is not rejected (p-value = 0.93). For reversals longer than 4 sec, the reversal duration is a good proxy for the total time of the manoeuvre. For reversals shorter than 4 sec, other effects such as variable turn times and simple turns (*n* = 1,276) should also be considered in determining the cost of the action. Kernel density estimation (KDE fit) is superimposed to aid visualization. b. The same plot as in a but excluding reversals from 4% gelatin clips (*n* = 340). P-value = 0.88. c. Density plot estimates (using KDE) for reversal counts per concentration show no systematic bias, hence allowing us to combine all reversals into a single distribution (in a, or with the exception of 4.0% gelatin, in b).

**Extended Data Figure 8:**
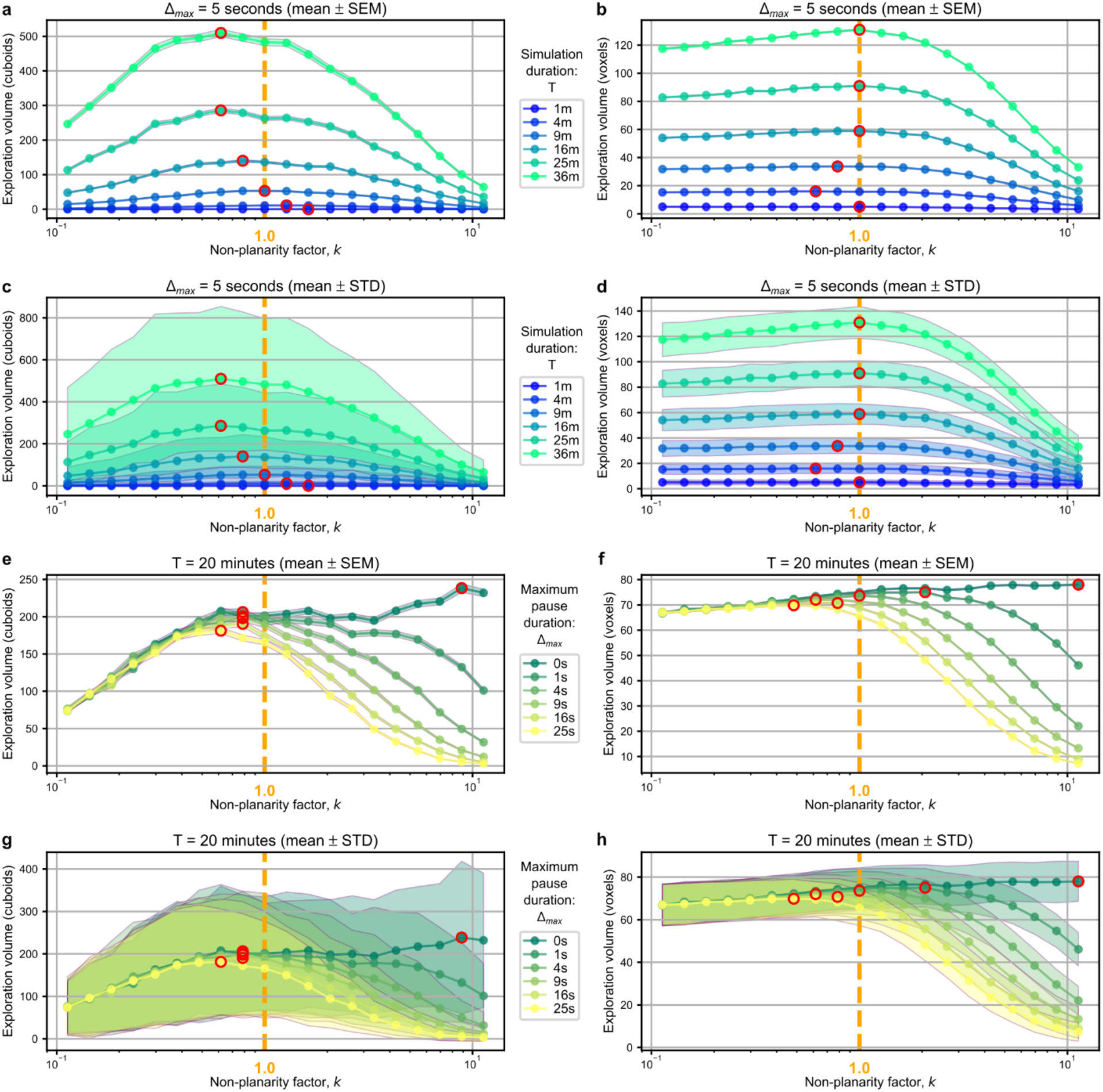
Optimality assessment of the model using alternative volume estimates. Exploration volumes as a function of non-planarity, *k* (as in Fig. 4e and f) reveal the largest exploration volumes (red circles). By definition, *k*=1 (orange dashed lines) corresponds to empirical data (Methods). Exploration volume is estimated as a cuboid with dimensions corresponding to the maximum distance travelled along each of the principal axes (a, c, e, g). Similar results are obtained when the volume is estimated by the number of unique voxels of size 0.5mm^3^ visited by the trajectory (For b, d, f, h). Standard deviations (STD) are high across instances (n=1,000 simulations for each combination of parameters) but standard error of the mean (SEM) is small. For both volume-estimation methods, the optimal strategy (i.e., optimal value of *k*) is robust to time-penalties that correspond to empirically observed reversal durations (2 ≤ Δ ≤ 20 sec, panels e-h) and local search durations (≤ 30 min, panels a-d). Hierarchical optimal strategies robustly appear in simulations with non-zero time-delay penalties for out-of-plane turns but are missing when Δ_max_ = 0 (e-h).

**Extended Data Table 1:** Run-and-tumble approximation. Pearson correlation coefficients between different run (n = 513) and turn (n = 565) metrics from the run-and-tumble approximation trajectories across the dataset (52 trajectories). Run durations and run speeds are normalised to account for variability in the gel concentrations and between animals. * p<0.05. The highly significant, strong negative correlation between run duration and speed motivates our use of a bivariate distribution in our model. The absence of significant correlations between θ and ϕ (|*R*_xy_| < 0.2) motivates the collapse of simple and complex turns into a single tumble state. We note that in our algorithm, any turns with small combined angles would not be registered as tumbles, explaining the co-absence of low θ, low ϕ tumbles. **†** All other statistically significant trends (p < 0.05) have negligible correlations (|*R_xy_*| < 0.2); turn angles are uncorrelated with run speeds or durations. Hence, these results justify a minimal two-state model (Fig. 4b, Methods).

| **x** | **y** | **Pearson Correlation Coefficient ($R_{xy}$)** | **p-value** |
| --- | --- | --- | --- |
| **Correlations between run statistics and turn angles motivate model parameterisation** |  |  |  |
| Run duration | Run speed | -5.265E-01 | **6.348E-38 \*** |
| In-plane turn angle ($\theta$) | Out-of-plane turn angle ($\phi$) | -1.634E-01 | **9.595E-05 \*** |
| **Correlations between run statistics and prior/subsequent turn angles** |  |  |  |
| Run duration | $\phi$ pre-run | -6.229E-02 | 1.589E-01 |
| Run duration | $\phi$ post-run | -9.636E-02 | **2.908E-02 †** |
| Run duration | $\theta$ pre-run | -4.025E-02 | 3.629E-01 |
| Run duration | $\theta$ post-run | -4.074E-02 | 3.571E-01 |
| Run speed | $\phi$ pre-run | 1.568E-01 | **3.626E-04 †** |
| Run speed | $\phi$ post-run | -1.487E-02 | 7.369E-01 |
| Run speed | $\theta$ pre-run | -5.719E-02 | 1.960E-01 |
| Run speed | $\theta$ post-run | -1.065E-01 | **1.579E-02 †** |
| $\phi$ pre-run | $\phi$ post-run | -4.290E-02 | 3.322E-01 |
| $\phi$ pre-run | $\theta$ post-run | -3.366E-03 | 9.394E-01 |
| $\theta$ pre-run | $\phi$ post-run | 6.491E-02 | 1.420E-01 |
| $\theta$ pre-run | $\theta$ post-run | 1.366E-01 | **1.922E-03 †** |
| Run duration pre-turn | Run duration post-turn | 4.130E-02 | 3.763E-01 |
| Run duration pre-turn | Run speed post-turn | -1.023E-01 | **2.815E-02 †** |
| Run speed pre-turn | Run duration post-turn | -1.042E-01 | **2.534E-02 †** |
| Run speed pre-turn | Run speed post-turn | 4.780E-02 | 3.057E-01 |
| **Correlations between pause statistics and turn angles** |  |  |  |
| Ratio of run spent paused | $\theta$ pre-run | 1.492E-02 | 8.529E-01 |
| Ratio of run spent paused | $\phi$ pre-run | 8.658E-02 | 2.809E-01 |
| Ratio of run spent paused | $\theta$ post-run | 6.916E-02 | 3.645E-01 |
| Ratio of run spent paused | $\phi$ post-run | -1.768E-01 | **1.962E-02 †** |
| Last pause duration | $\phi$ post-run | -1.079E-01 | 2.954E-01 |
| Last pause duration | $\theta$ post-run | 3.152E-02 | 7.604E-01 |

